# Kernels of Motor Memory Formation: Temporal Generalization in Bimanual Adaptation

**DOI:** 10.1101/2024.01.03.574029

**Authors:** Ian S. Howard, Sae Franklin, David W. Franklin

**Affiliations:** School of Engineering, Computing and Mathematics, University of Plymouth, Plymouth, United Kingdom; Neuromuscular Diagnostics, Department Health and Sport Sciences, TUM School of Medicine and Health, Technical University of Munich, Germany; Munich Institute of Robotics and Machine Intelligence (MIRMI), Technical University of Munich, Munich, Germany; Munich Data Science Institute (MDSI), Technical University of Munich, Munich, Germany

**Keywords:** motor learning, bimanual context, temporal generalization, dynamic learning, human, viscous curl field

## Abstract

In daily life, we coordinate both simultaneous and sequential bimanual movements to manipulate objects. Our ability to rapidly account for different object dynamics suggests there are neural mechanisms to quickly deal with them. Here we investigate how actions of one arm can serve as a contextual cue for the other arm, and facilitate adaptation. Specifically, we examine the temporal characteristics that underlie motor memory formation and recall, by testing the contextual effects of prior, simultaneous, and post contralateral arm movements in both male and female human participants. To do so, we measure their temporal generalization in three bimanual interference tasks. Importantly, the timing context of the learned action plays a pivotal role in the temporal generalization. While motor memories trained with post adaptation contextual movements generalize broadly, motor memories trained with prior contextual movements exhibit limited generalization, and motor memories trained with simultaneous contextual movements do not generalize to prior or post contextual timings. This highlights temporal tuning in sensorimotor plasticity: different training conditions yield substantially different temporal generalization characteristics. Since these generalizations extend far beyond any variability in training times, we suggest that the observed differences may stem from inherent differences in the use of prior, current and post-adaptation contextual information in the generation of natural behavior. This would imply differences in the underlying neural circuitry involved in learning and executing the corresponding coordinated bimanual movements.

**Significance Statement:** This study addresses a fundamental question in the field of sensorimotor neuroscience of how multiple movements are temporally linked within a single motor memory. We examine the temporal generalization of motor memory formation by varying the timing of contextual movements associated with a learned motor memory across a range of prior, current, and post adaptation movement times using a bimanual motor learning task. We observed distinct patterns of temporal generalization based on whether the contextual movements occurred prior to, simultaneously with, or after the adaptation movement. For the first time, our findings reveal that the timing of contextual movements is crucial in the formation and generalization of motor memories.

## Introduction

Many everyday tasks require coordinated actions of our two arms. While some tasks, such as opening a jar, require the simultaneous control of both arms, others, such as juggling, involve an initial movement of one hand followed by a subsequent movement of the other hand. Although juggling involves the manipulation of objects that vary in size, weight, or shape, professional jugglers demonstrate impressive hand-eye coordination, producing rapid and accurate object trajectories. Such effective manipulation requires that our sensorimotor system accurately accounts for the dynamics of the objects currently being manipulated, as well as those to be manipulated in the future. Motor learning of such object dynamics during bimanual movement is therefore a fundamental process underpinning human skills.

Adaptation to dynamics occurs through the formation of a motor memory (Lackner and Dizio, 1994; Shadmehr and Mussa-Ivaldi, 1994; Kawato, 1999; Shadmehr et al., 2010). However, these motor memories are not limited to the specific tasks learned, but they generalize to similar movements and tasks, allowing flexible and adaptable motor control (Conditt et al., 1997; Thoroughman and Shadmehr, 2000; Donchin et al., 2003; Orschiedt and Franklin, 2023). As this generalization is thought to stem from learning via neural basis functions, measurement of the extent and type of generalization allows for the investigation of the tuning of these neural basis functions within the sensorimotor control system. Here we use the term temporal kernel to describe the temporal tuning of the generalization, which may arise from the underlying neural basis functions.

To study the formation, selection and generalization of motor memories and the effect of context, two opposing force fields are often randomly presented during reaching movements. Without any additional cues, this random presentation results in interference: learning one force field interferes with the memory of the other (Shadmehr and Brashers-Krug, 1997; Karniel and Mussa-Ivaldi, 2002; Osu et al., 2004; Krakauer and Shadmehr, 2006). However, the association of the force field with certain contextual cues allows learning of the two independent motor memories and appropriate switching of the memories, termed dual adaptation (Nozaki et al., 2006; Hirashima and Nozaki, 2012; Howard et al., 2012). This paradigm allows us to examine which contextual cue types can select and switch between different motor memories within the sensorimotor control system, and identify those cues that do not have such functions (Howard et al., 2013; Forano et al., 2021).

While the tuning of neural basis functions that underly generalization is typically examined as a function of both adaptation and contextual movement directions (Thoroughman and Shadmehr, 2000; Donchin et al., 2003; Howard and Franklin, 2015, 2016; Sarwary et al., 2015), and of contextual movement kinematics (Howard et al., 2020; Orschiedt and Franklin, 2023), the temporal generalization of these motor memories, examined by varying the timing of contextual movements, has yet to be investigated.

Here we use an interference task to examine how different temporal relationships between motion of the two hands affect adaptation, specifically focusing on prior, simultaneous, and post contextual motions. In unimanual tasks, both prior contextual lead-in movements and post adaptation contextual follow-through movements allow dual adaptation (Howard et al., 2012, 2015; Sheahan et al., 2016). While in bimanual tasks, simultaneous (Nozaki et al., 2006; Howard et al., 2010) and prior (Gippert et al., 2023) contextual movements support dual adaptation, little is known about the role of post adaptation contextual movements. Moreover, we have no measure of the temporal generalization of these motor memories: how learning at one timing relation transfers to other timing relations. Here, we first investigate how prior, simultaneous and post adaptation contextual movements of the contralateral hand affect learning opposing force fields. For each trained timing relationship, we then examine the recall of this learned motor memory to a range of different temporal relations, enabling us to map out the temporal generalization of these motor memories across different temporal relations, and extract the temporal kernels of motor memory formation and recall.

## Methods

### Experimental Setup

#### Participants

Twenty-six human participants were randomly allocated to three experiments. Two participants were excluded from further analysis purely based on their adaptation to the curl force field, leaving twenty-four. Specifically, one participant was excluded as they did not adapt at all to the viscous curl force field (final adaptation level at the end of day 2 < 10%). The other participant was removed as the kinematic error reduced to below 5 mm within the first 100 trials in the force field while force compensation remained variable and low throughout the experiment, suggesting either co-contraction or more cognitive strategies involved in adaptation. The generalization data of these two participants was not analyzed. This resulted in eight participants for each of Experiment 1 (3 females, average age 24.6 ± 2.4 years), Experiment 2 (5 females, average age 28.4 ± 8.5 years), and Experiment 3 (3 females, average age 25.6 ± 2.7 years). All participants were right-handed as determined by the Edinburgh Handedness Questionnaire (Oldfield, 1971) and naïve to the purposes of the experiment. Prior to the experiment, each participant provided written informed consent. The experiments were conducted for Plymouth participants (n=3), in accordance with ethics approval by the University of Plymouth ethics committee and conducted for Munich participants (n=23) in accordance with ethics approval by the Ethics Committee of the Medical faculty of the Technical University of Munich. Each participant completed one experiment, consisting of two three–hour experimental sessions over two consecutive days, totaling approximately six hours for each participant.

#### Apparatus

Participants performed all experiments using a bimanual vBOT planar robotic manipulandum paired with an associated virtual reality system (Howard et al., 2009b) (Fig. 1A). Handle positions were calculated based on measures using optical motor encoders sampled at 1000 Hz, and torque-controlled motors applied end-point forces. A force transducer (Nano 25; ATI), mounted under the left handle, measured the applied forces. These force signals were low-pass filtered at 500 Hz using 4th-order analogue Bessel filters before digitization. To minimize body movement, participants were comfortably seated in a sturdy chair positioned in front of the apparatus and were firmly strapped against the backrest with a four-point seatbelt.

**Figure 1.**
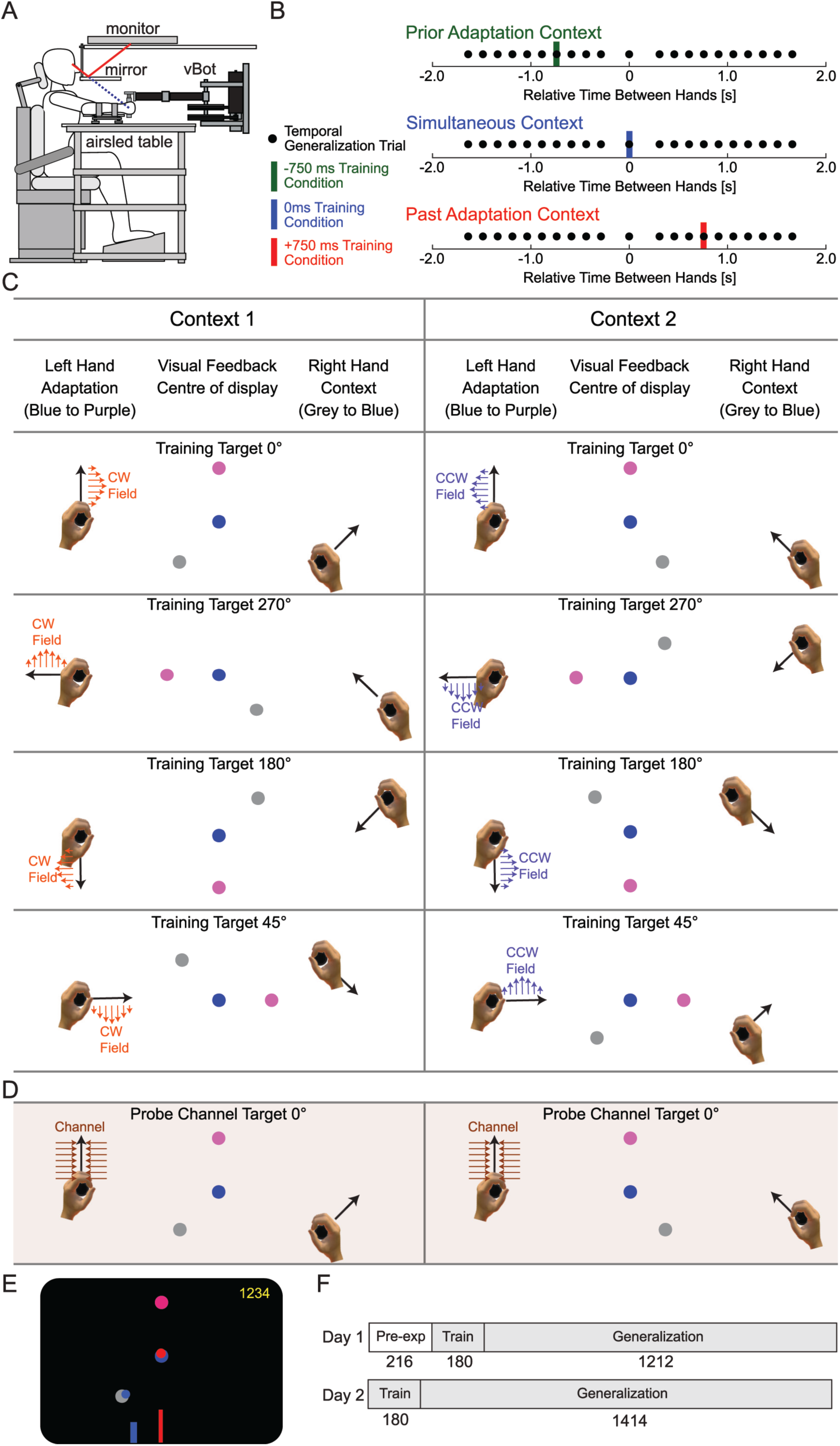
Experimental Setup and Design. **A.** Experiments were conducted using a bimanual vBOT setup in a virtual reality environment. The participant sits while grasping both handles of the vBOT manipulandum and receiving visual feedback in the plane of movement. **B.** Bimanual Movement Timing Relationships. Three separate experiments were conducted with training at timing relationships of +750 ms (Post Adaptation Context; red), 0 ms (Simultaneous Context; blue), and -750 ms (Prior Context; green). The training time relation between the two hands is shown with the colored bar for each experiment. To test temporal generalization, channel trials were presented at 21 distinct timing relationships, represented by black circles, ranging from - 1.8 to +1.8 seconds. Each temporal relationship was tested seven times for each of the two contexts, amounting to a total of 14 tests per relationship. **C.** Experimental protocol for training and testing. The top four rows depict the contextual relationships used in both null and training trials for the four possible adaptation movement directions. The movement of the right-hand provides context, with four distinct starting locations (at 45°, 135°, 225°, and 315°) leading to a central location on the right. The left-hand performs the adaptation movement, executed either in a null field or a viscous curl field. This hand also has four movement directions, beginning from the central position on the left and progressing to specific target locations (0°, 90°, 180°, and 270°). Notably, each adaptation movement direction is paired with two unique contextual movement directions (two columns indicating the two possible contexts). It can be seen that while the two hands were separated by 30 cm, the visual representations of the two hands’ movements were aligned such that the end of the movement of the right-hand ended in the center target where the left-hand starts its movement. This involved a visual shift of 15 cm for each hand. **D.** The row (brown shaded region) displays the contextual relationships for channel trials, designed to estimate feedforward adaptation to the viscous curl field. These channel trials proceed solely along the y-axis to an upper target and are linked with two distinct angular contextual movements originating at 135° (right column) and 225° (left column). **E.** Visual feedback provided to the participants. At the bottom of the screen the temporal relation of the right-hand (blue bar) and left-hand (red bar) for the next trial was displayed. The red bar was always in the center of the screen but the blue bar was moved depending on the timing of the relation. Cursors representing the right (blue) and left (red) hands were shown, along with the start and end targets for both movements. The red cursor was moved to the red target while the blue cursor was moved to the blue target. A score representing the number of successful targets was presented at the top right of the screen. **F.** The experimental trial schedule was identical across all three experiments, but each experiment used different training timing relationships. Day 1 started with pre-exposure (Pre-exp) in the null field trials which included testing probe channel trials. The white color indicates that all non-channel trials were performed in a null field to familiarize participants with the manipulanda and the testing procedures. The curl force fields (grey shaded region) were introduced in the Train session (180 trials), followed by a Generalization phase (1212 trials) that provided practice to the bimanual movement relations. Day 2 started with further training (Train), followed by a final Generalization session to study generalization.

During the experiment, participants held the left and right robot handles with their respective hands, with their forearms supported by air sleds. This setup effectively restricted arm movements to the horizontal plane and provided full support of the mass of their arms. Participants could not directly observe their arms or hands; instead, veridical visual feedback was provided using a 2D virtual reality system to overlay images of the left- and right-hand starting and target locations (1.25 cm radius) and hand cursors (a 0.5 cm radius). This arrangement ensured that the visual cursor appeared to the participant in the same plane as their hand. Data were sampled at 1000 Hz and stored for offline analysis.

#### Force Fields

Right-hand movements, termed contextual movements, were always performed in a null field. Left-hand movements, termed adaptation movements, were performed in one of three environments: null field, curl force field, or channel trials. In the null field trials, no external forces were applied on the hands by the robotic manipulandum. In the curl force field trials, a velocity-dependent curl force field (Gandolfo et al., 1996) was implemented as:

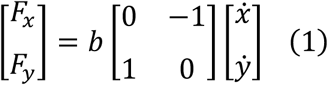

The viscous field constant b was set to ±13 Nm^-1^s, with the sign determining the force field’s direction, which was either clockwise (CW) or counter clockwise (CCW). Each participant experienced both force field directions, where the direction of the force field was consistently associated with specific contextual movements of the participant’s right-hand. The relationship between the directions of the contextual movements and the curl field direction (CW/CCW) was counterbalanced across participants. In channel trials, a mechanical channel, from the movement start position to the center of the final target, was applied perpendicular to the motion direction (Scheidt et al., 2000; Milner and Franklin, 2005) with a stiffness of 6,000 Nm^-1^ and a damping of 30 Nm^-1^s. More specifically, the channel was initiated at the time of the go cue and was oriented from the participant’s hand position at the time of the go cue to the center of the final target. Channel trials were only conducted for movements towards the 0° target.

### Experiments

We conducted three bimanual interference experiments, each involving a different participant group, to investigate the temporal generalization of contralateral arm movements. Each experiment trained participants at a specific timing relation between the two hands and then assessed their ability to generalize this training across a broad range of timing relations (Fig. 1B). The main difference between experiments was the bimanual timing relationship for the training movements, which is the temporal gap between contextual and adaptation movements (Fig. 1B, colored bars). Almost all other aspects of the three experiments were identical.

In all experiments, the right-hand (contextual movement) moved from a start location to the center point, while the left-hand (adaptation movement) moved from the center point to a target location. All trials involved 12 cm active movements of both hands. The bimanual timing relation of the training movements was different for each experiment (Fig. 1B).

#### Experiment 1 – Post Adaptation Context

Participants first executed the left-hand adaptation movement, followed by the right-hand contextual movement. The adaptation movement starts 750 ms before the contextual movement, and is defined as a bimanual timing of +750 ms.

#### Experiment 2 – Simultaneous Context

Participants performed both the contextual and adaptation hand movements simultaneously, corresponding to a bimanual movement timing of 0 ms.

#### Experiment 3 – Prior Context

Participants first executed the right-hand contextual movement, followed by the left-hand adaptation movement. The contextual movement starts 750 ms before the adaptation movement. This timing is defined as -750 ms.

### Experimental Design

The left-hand adaptation movement was associated with either a clockwise (CW) or counter clockwise (CCW) curl force field during the exposure phase of the experiment. Adaptation movements were oriented at 0°, 90°, 180° and 270°, each paired with two contextual movements rotated at either 45° CW or 45° CCW, to provide the contextual cue (see Fig. 1C). This resulted in eight unique combinations of contextual and adaptation movements. For half of the participants, context 1 was associated with the CW curl force field and context 2 was associated with the CCW curl force field while this was reversed for the remaining participants. Although the physical locations of the handles were shifted away from the center of the workspace to avoid hand collisions, the visual feedback for the task was centered in the workspace, aligning the contextual movement endpoint with the adaptation movement start. Right-hand contextual movements always occurred in null fields. During the exposure phase, the left-hand adaptation movements were always performed in either a curl force field or a mechanical channel.

All trials consisted of a two-part movement produced by both hands. Each trial began when the two cursors, representing the two hands, were placed in their respective starting circles. Participants could not see their arms and hands, but the hand cursors were color coded: red for the left cursor and blue for the right cursor. The starting circle for the right-hand was grey, while that for the left-hand was blue. Participants were instructed to keep their hands in these circles until an auditory cue signaled the start of the movements. The order of the hand movements was displayed at the bottom of the screen, with red and blue bars indicating the relative timing of each hand’s movements on the time scale along the x-axis (Fig. 1E), which prepares the participant to move the hands with a specific order and relative timing. The exact timing of the movements for both the right- and left-hands were then cued in a trial with auditory beeps.

Upon the presentation of each auditory cue, participants moved each of their hands (order as indicated by the display at the bottom of the screen) to the appropriate target circle and stopped. Once the red cursor (left-hand) was successfully placed into the final target circle, the circle’s color changed from pink to red. Similarly, when the blue cursor (right-hand) reached the center target circle, the circle’s color changed from light to dark blue. Importantly, visual feedback of the trajectory of the contextual movement was never shown to the participants due to its strong contextual effect (Howard et al., 2012, 2020; Howard and Franklin, 2016; Franklin et al., 2023). Instead, participants relied on this color change to confirm successful cursor placement in the target circle.

In each experiment, the time intervals between the start of the movement for the left- and right-hands were set at +750 ms, 0 ms, or -750 ms for the training trials. The trial ended once both hands remained stationary within their respective target locations. At the end of each trial, the robotic manipulandum passively moved the participants’ hands to the next starting locations. The next trial began once the handles had been stationary within the start locations for 300 ms. Participants were required to take short breaks approximately every 200 trials but could also rest at any time between trials by moving the cursors out of the start targets.

At the end of each trial, participants received feedback regarding the peak movement velocity of each hand. This speed feedback appeared immediately after every trial as two vertical bars displayed at the top left of the screen, each representing a hand. The desired speed range was indicated by two horizontal lines corresponding to 45.0 cm/s ± 15%. If a hand’s peak speed was within this range, its bar turned green. Conversely, if the movement was too fast or too slow, the bar turned red. Participants were encouraged to maintain their peak movement speed within this range.

If a participant’s hand moved from the start position before the auditory cue, or if they failed to complete a trial by moving their hands to the targets within the maximum time limit (5 seconds), the trial was cancelled. Trials were also cancelled if the time difference between initiating the movements of both arms exceeded 200 ms for simultaneous context experiments and 950 ms for the prior and post context experiments. Cancelled trials were repeated at the end of each block. A trial was considered successful if both hands moved with a peak speed within the specified range. For each successful trial, one point was added to the participant’s score, displayed at the top right of the screen.

#### Probe Channel Trials

To estimate both the predictive compensation for the force field as part of the adaptation process, and to measure the temporal generalization across a range of timing relations between the two limbs, we introduced probe channel trials. Throughout the experiment, on randomly selected trials, channel trials matching the training condition were introduced to estimate the feedforward adaptation to the curl force field. These channel trials only occurred on adaptation movements (left-hand) moving forward to the final target, and were associated with two possible angular contextual movements (as shown in Fig. 1D).

To examine the generalization of learning related to contextual movements made at specific temporal training relationships, we assessed the recall of adaptation across a range of timing relationships. Specifically, channel trials were conducted at 21 different bimanual timing relationships: [-1.8, -1.65, -1.5, -1.35, -1.2, -1.05, -0.9, -0.75, -0.6, -0.45, 0, 0.45, 0.6, 0.75, 0.9, 1.05, 1.2, 1.35, 1.5, 1.65, 1.8] seconds. The bimanual timing relationships used in the three experiments, as well as those used to probe generalization, are illustrated in Fig. 1B (black dots). All probe channel trials were limited to the forward reaching movements (target 0°) only.

### Protocol

Participants performed two experimental sessions on two consecutive days. On day 1, participants completed 1608 trials, and on day 2, they completed 1594 trials, totaling 3202 trials (Fig. 1F). Temporal generalization probe trials were performed on day 1 in both pre-exposure and generalization phases to familiarize participants with executing movements of the two arms at different timing relations.

#### Day 1

##### Pre-exposure phase

216 trials. Participants began with 36 trials in the null field, divided into two blocks of 18 trials each. Each block consisted of sixteen null field trials (two for every movement type) and two channel trials (one for each context). This was followed by two blocks of mixed null field and generalization trials, totaling 180 trials. Each of these blocks comprised 48 null field trials and 42 generalization channel probe trials (1 for each context at each of the 21 different timing relationships).

##### Training phase

180 trials. Participants performed ten blocks of trials in the curl force fields, where each curl force field was associated with one of the two contextual cues. Each block, consisting of 18 trials, included 16 curl field trials (two for every movement) and 2 channel trials (one for each context).

##### Generalization phase

1212 trials. Participants performed six blocks in the curl force fields. Each block was composed of 160 curl field trials (20 for each of the 8 possible movements) and 42 generalization channel probe trials (one for each of the 21 different timing relations for both contexts).

#### Day 2

##### Training phase

180 trials. Participants performed ten blocks where each block comprised 16 curl field trials and 2 channel trials.

##### Generalization phase

1414 trials. Participants performed seven blocks. Each block consisted of 160 curl field trials and 42 generalization channel probe trials (1 for each of the 21 different timing relations for both contexts). Data from this phase was used to measure the temporal generalization.

### Data Analysis

The analysis was conducted offline using MATLAB R2023a. To examine learning and generalization, kinematic error and force compensation were calculated for each trial.

#### Kinematic Error

For each trial performed in null and viscous curl force fields, the kinematic error was determined in the adaptation phase of the movement by calculating the signed Maximum Perpendicular Error (MPE). The MPE is defined as the maximum deviation of the hand’s path from a straight line between the starting location and the target. For each participant, the curl-field direction dependent average MPE was computed over a set of 8 trials. In addition, an average MPE was computed over both curl-field directions over a set of sixteen trials for each participant. To combine results from Clockwise (CW) and Counterclockwise (CCW) field trials, the sign of the MPE was appropriately adjusted. Subsequently, the mean and standard error (SE) of the MPE were calculated across all participants.

#### Force Compensation

During each channel trial, the force exerted by the participants perpendicular to the wall of the simulated channel was measured to estimate predictive feedforward adaptation. This measure was used to assess both learning and generalization to different temporal relations between the two hands. Force compensation was calculated by regressing the measured force on the channel wall against the force required to perfectly compensate for the force field (calculated as the movement velocity multiplied by the force field strength b) (Smith et al., 2006). No offset term was used in the fitting procedure. This method offers greater accuracy than relying solely on a reduction in kinematic error, as it avoids the effect of limb impedance due to muscle co-contraction (Franklin et al., 2003; Milner and Franklin, 2005; Franklin and Franklin, 2021, 2023). Additionally, it provides an estimate of the adaptation throughout the entire movement, rather than at just a single time point. To examine adaptation, the sign of the force compensation values for the two different contexts were maintained, so that adaptation to both field directions could be observed independently. In additional analysis, the sign-corrected compensation values were averaged over the two adjacent channel trials within each block of 18 trials, representing the two different contexts. In both cases, the mean and SE of these force compensation values were calculated and analyzed across all participants throughout the experiment.

#### Bimanual Timing Relation

Channel probe trials examined the generalization of force compensation across 21 different timing relations between the limbs, spanning a ±1.8 second range. Although participants were cued to initiate movements with these specific timing relations, they often moved with slightly different timing relations. To estimate the actual bimanual timing relations executed by participants, we calculated the start of each movement based on specific position and speed criteria. A hand was considered to have initiated movement when it deviated more than 2 mm from the starting target radius (1.27 cm from the center of the start) and was traveling at a speed greater than 5 cm/s. A movement was considered to have terminated when it reached within 2 mm from the target radius (1.27 cm from the center). The bimanual temporal relation for a given trial was calculated by subtracting the start time of the left (adaptation) hand from the start time of the right (contextual) hand. Negative values indicate that the contextual movement was made before the adaptation movement.

#### Temporal Generalization

To quantify the temporal generalization of the motor memory, we used force compensation data from the channel trials during the Generalization Phase on the second day (refer to Fig. 1E) as participants had achieved a higher level of adaptation. Temporal generalization was quantified based on the actual timing relations between the two hands. For each participant, compensation estimates from the channel trials were first allocated to 29 temporal bins based on their bimanual timings. The center of each bin ranged from -2.1 to +2.1 seconds, spaced by 150 ms intervals, matching the cue timings (and widths) used in the generalization probe channel trials.

For each bin, we calculated the mean and the number of bin counts for each participant. Bins with no data for a participant were removed to avoid misleading estimates. To estimate generalization as a function of temporal relationships, we calculated the mean and standard error of the mean (SEM) for the bin values across all eight participants.

### Statistics

Statistical analyses were performed in JASP 0.18.1 (JASP Team, 2023). A Mauchly test was applied to check for sphericity where applicable. In the case that the Mauchly test was significant, the degrees of freedom were adjusted using a Greenhouse-Geisser correction. The *α*-level was set to .05 in all tests, and omega squared (ω^2^) was reported as the effect size. Omega squared is an estimate of how much variance in the measurement can be accounted for the explanatory variables.

For each experiment, we performed repeated measures ANOVAs to examine differences in the MPE and force compensation within each experiment. We compared the final pre-exposure MPE (mean of last block in pre-exposure), initial exposure MPE (mean of first block in exposure) and final exposure MPE (mean of final block on day2) using a repeated measures ANOVA with repeated factor of phase (3 levels). If a significant main effect was found, post-hoc comparisons were performed (Holm-Bonferroni corrected). For the force compensation, we compared the final pre-exposure (mean of last block in pre-exposure) and final exposure (mean of final block on day2) using a repeated measures ANOVA with a repeated factor of phase (2 levels: pre-exposure and final exposure).

To compare across experiments, we performed ANOVAs on the MPE and force compensation in the final exposure block with a factor of experiment condition (3 levels). If a significant main effect was found, post-hoc comparisons were performed (Holm-Bonferroni corrected).

To compare the temporal generalization within each experiment and across the three experiments we performed a repeated measures ANOVA with a repeated factor of timing (21 levels) and between subjects factor of experimental condition (3 levels). If a significant main effect was found, post-hoc comparisons were performed (Holm-Bonferroni corrected for the whole family of comparisons). Only post-hoc comparisons which were relevant for the study were examined and reported. Specifically, we only examined differences across timings within one experiment, or differences across experiments for each timing. To support these findings without binning the data or averaging trials which reduces the variability of the data, we also performed statistical analysis on the full data set (all trials from all participants) using a Generalized Additive Mixed Model (GAMM) in R (4.4.1). We fit the model (force compensation as a function of time; random effect of participants) both with conditions (3 experiments; full model) and without conditions (reduced model) as a factor. We then compare the two models (full and reduced models) with a likelihood ratio test to determine which model has a significantly better fit. To perform pairwise comparisons between experimental conditions, we use a similar procedure with each pairwise comparison of experiments. For each comparison, we fit both the full model with two conditions and the model with no conditions. Again, a likelihood ratio test was then used to test which model best explains the data. If the model with conditions better explains the data, this indicates that there is a significant effect of condition.

## Results

Three groups of participants performed separate experiments, with each group experiencing a different timing relationship in bimanual training while adapting to two opposing force fields (Fig. 1B, C). Each group tested whether prior, simultaneous, or post contralateral arm movements can provide a contextual cue for the ipsilateral arm. Here the ipsilateral arm was exposed to opposing viscous curl fields where the direction of the field was determined by the contralateral arm motion. After this adaptation phase, the generalization of the learned motor memory was assessed using probe channel trials (Fig. 1D) across various timing intervals between the two arms. These intervals included timings before (prior context), during (simultaneous context), and after (post context) the adaptation movement (Fig. 1B).

The generalization was examined across the three conditions and the different timings using a repeated measures ANOVA finding a significant main effect of timing (Greenhouse-Geisser corrected: F_3.204,67.279_=5.863; p=0.001; ω^2^=0.131), between subjects’ effect of experiment condition (F,_2,21_=12.493; p<0.001; ω^2^=0.258), and interaction between timing and experimental condition (Greenhouse-Geisser corrected: F_6.407,67.279_=23.296; p<0.001; ω^2^=0.544). Post-hoc comparisons (Holm-Bonferroni corrected) of the interaction effects were performed and reported in subsequent sections.

### Experiment 1: Post Adaptation Context

Experiment 1 (post adaptation context) tested the training condition of +750 ms, in which the right-hand contextual movements were performed 750 ms after a preceding left-hand adaptation movement. That is, the movement of the right-hand acts as a planned follow-through movement that takes place after the adaptation movement of the left-hand. Before the introduction of the force field, participants made straight movements to each of the four targets with their left-hand (Fig. 2A). Once the force field was introduced, participants’ movements were strongly curved away from the straight line, with the disturbance direction depending on the sign of the force field (Fig. 2B). However, by the end of the exposure phase the movements were disturbed less laterally, and participants made straighter movements to the targets for both force field directions suggesting adaptation to the dynamics (Fig. 2C).

**Figure 2:**
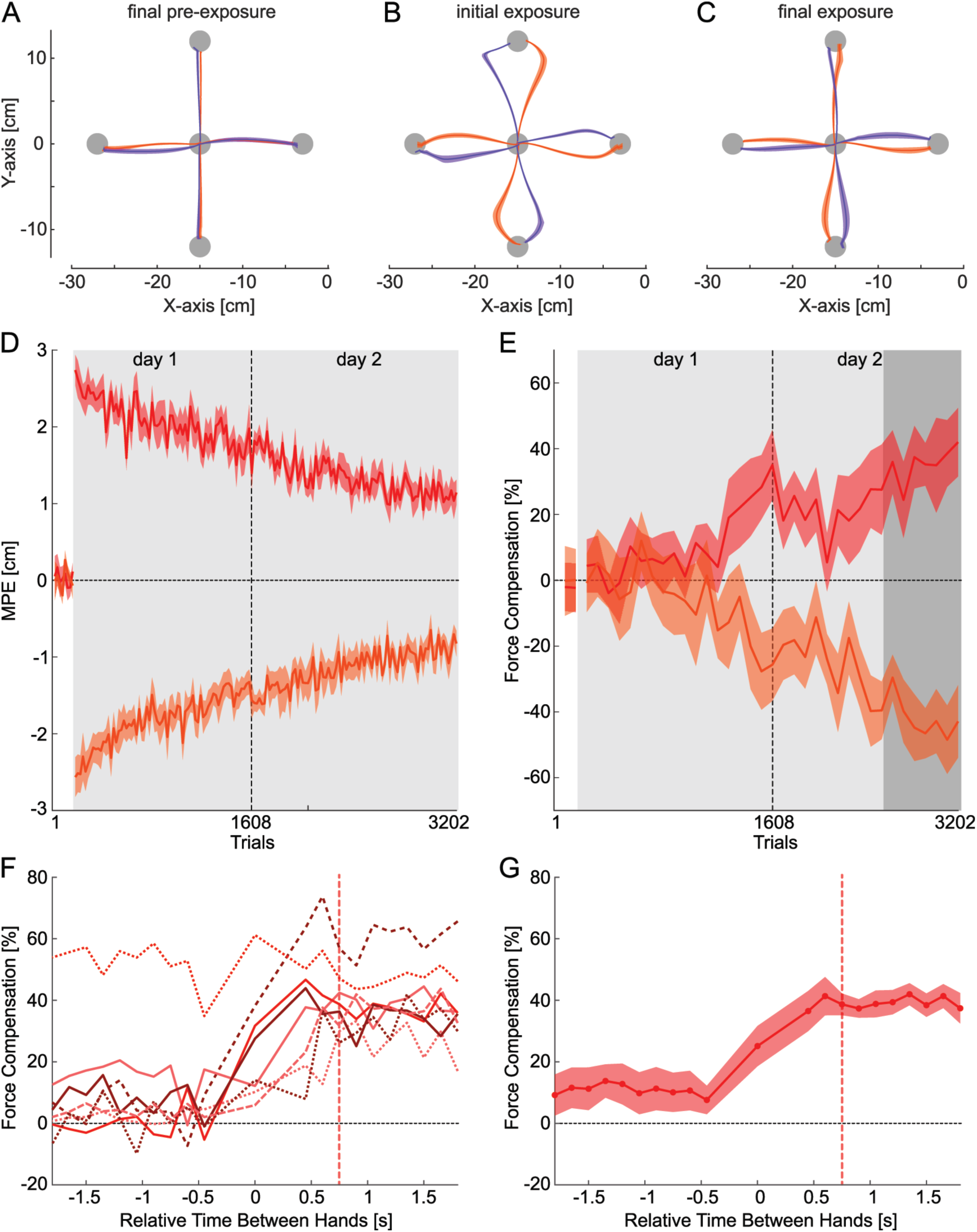
Results for Experiment 1 where the right-hand contextual movement follows the left-hand training movement by 750 ms (+750 ms condition). **A.** Hand paths in the last block of pre-exposure in the null field condition. The mean and standard error (shaded region) of left-hand paths across participants from the central location to the target are shown for both contextual cue movement directions. Movements that will be associated with the CW curl force field are shown in orange whereas movements that will be associated with the CCW force field are shown in purple. Within the selected block, the last 8 unique movement directions per participant were used for the calculation. **B.** The hand paths on initial exposure to the curl field. **C.** The hand paths in the final block of curl field exposure. **D.** The kinematic error, measured as maximum perpendicular error (MPE) for the left-hand, plotted over the course of the experiment. The mean (line) and standard error (shaded area) are calculated across the participants for each curl force field direction separately. The grey shaded region indicates the period where the force field is applied to the left-hand during the adaptation movements. **E.** Force compensation, expressed as a percentage of complete adaptation to the force field, over the course of the experiment for each curl force field direction separately. Positive values indicate adaptation to CW field whereas negative values indicate adaptation to the CCW field. Grey region indicates the application of the force field, and the dark grey region indicates the period over which the generalization to different timing relations between the two hands was measured. **F.** The temporal generalization of the learned motor memory to all timing relations between the two hands shown for individual participants. Each line denotes a different participant, and the red dashed vertical line indicates the trained relation. **G.** The mean and standard error of the temporal generalization of the learned motor memory across participants. Red circles denote the center of the window over which values were averaged. We see that training at +750 ms led to effective generalization to post adaptation contextual movement timing relationships, but not to prior adaptation contextual movements.

To quantify adaptation, we examined two measures: maximum perpendicular error (MPE) and force compensation. A repeated measures ANOVA indicated that MPE showed significant variations between the pre-exposure, initial exposure and final exposure phases (F_2,14_=129.514; p<0.001; ω^2^=0.864), which were further tested with post-hoc comparisons. MPE (Fig. 2D) was close to zero in the pre-exposure phase, but increased immediately upon the introduction of the force field (p<0.001). MPE then reduced over the course of the experiment, with final levels smaller than during initial exposure (p<0.001) but not as low as in pre-exposure (p<0.001). Force compensation, as measured in channel trials for the training timing relation, was close to zero in the pre-exposure phase. However, force compensation increased slowly during the exposure phase, reaching around 40% by the end of the second day (Fig. 2E). A repeated measures ANOVA indicated a significant increase in force compensation between final pre-exposure and final exposure levels (F_1,7_=188.732; p<0.001; ω^2^=0.887).

The temporal generalization of the learned motor memory was examined towards the end of the exposure phase by introducing probe channel trials. In these trials, the timing between the right contextual hand movement and the left-hand adaptation movement was changed from +750 ms to one of 21 values between -1.8 and +1.8 seconds (Fig. 1B). The force compensation observed in these trials enabled us to assess the extent to which the learned motor memory was specific to the trained condition, or whether it generalized to other timing relations. These timing relations included both those matching the trained relation between the hands (prior adaptation contextual movements) and extended to those that did not (synchronous or post adaptation contextual movements).

The temporal generalization of the motor memory after training at +750 ms demonstrates consistent and effective generalization to other post adaptation contextual movements (Fig. 2G), with all timing relations between +1.8 and +0.45 s exhibiting around 40% compensation. However, there was little transfer to adaptation from prior contextual movements (approximately 10%). Post-hoc comparisons supported these findings with no significant differences across any post movement timings (all p=1.0) but significant differences between all post movements and all prior movements (all p<0.001), and no significant differences between all post movement timings (all p=1.0). While the simultaneous condition was different than the nearest prior condition (p=0.041), there were no significant differences with any of the other post (all p>0.078) or prior conditions (all p>0.188). This generalization pattern was consistent across all but one participant, who showed flat generalization across all training relations (Fig. 2F). We believe this flat generalization may indicate a more explicit adaptation occurred in this participant, although we have no evidence to test this possibility directly. Overall, we find that the post contextual bimanual movement generalizes strongly to all other post contextual movement timings and partially to the synchronous condition. However, there is little or no generalization to the prior adaptation movement contextual conditions.

### Experiment 2: Simultaneous Context

In the second experiment (simultaneous context), the onset of both the adaptation and contextual movements occurred simultaneously (0 ms). In the pre-exposure phase, participants made straight movements towards each of the four targets with their left-hand (Fig. 3A). Again, once the force field was introduced, participants’ movements were strongly disturbed in the direction of the applied force field (Fig. 3B). However, by the end of the exposure phase, the participants made straight movements to the targets for both force field directions (Fig. 3C). The kinematic error (MPE) increased immediately upon introduction of the force field (Fig. 3D), but reduced over the course of force field exposure. A repeated measures ANOVA found significant differences across the phases of the experiments (F_2,14_=105.044; p<0.001; ω^2^=0.886), with initial exposure larger than pre-exposure (p<0.001) and final exposure (p<0.001), but no difference between pre-exposure and final exposure (p=0.097). Force compensation steadily increased throughout the exposure phase, reaching approximately 50% by the end of the second day (Fig. 3E). A repeated measures ANOVA found a significant increase in force compensation between final pre-exposure and final exposure levels (F_1,7_=117.522; p<0.001; ω^2^=0.867). Thus, participants were able to adapt to the two opposing force fields with simultaneous contextual movements of the other hand.

**Figure 3:**
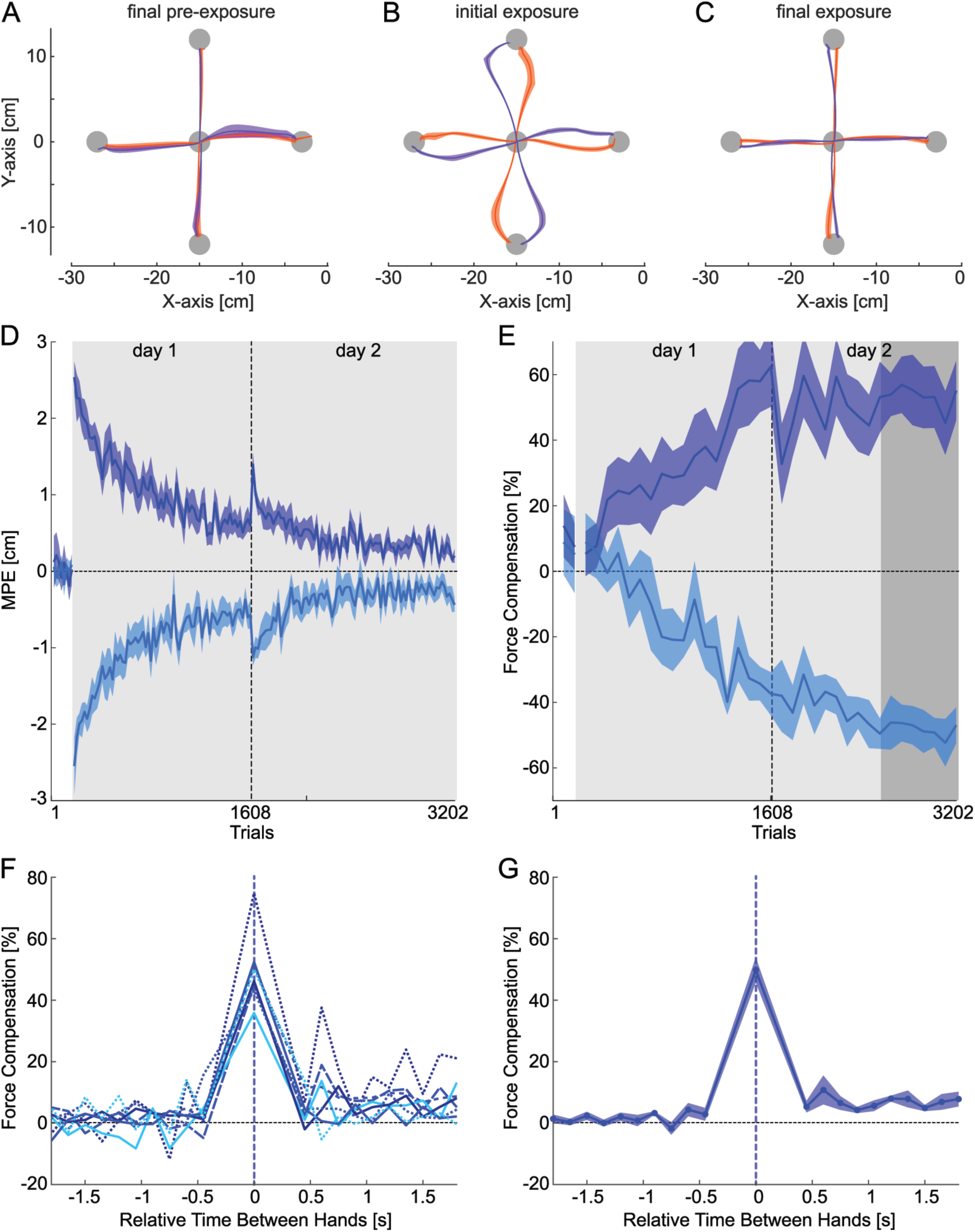
Results for Experiment 2 using training at a 0 ms synchronous context timing relationship **A.** Hand paths in the last block of pre-exposure in the null field condition are shown for both contextual cues. **B.** The hand paths on initial exposure to the curl field. **C.** The hand paths in the final block of curl field exposure. **D.** The kinematic error, measured as maximum perpendicular error (MPE) for the left-hand, plotted over the experiment for each curl force field direction separately. **E.** Force compensation for each curl force field direction separately, expressed as a percentage of complete adaptation to the force field, over the experiment. Grey region indicates the application of the force field, and the dark grey region indicates the period over which the generalization to different timing relations between the two hands was measured. **F.** The temporal generalization of the learned motor memory to all timing relations between the two hands, shown for individual participants. Each line denotes a different participant and the blue dashed vertical line indicates the trained relation. **G.** The mean and standard error of the temporal generalization of the learned motor memory across participants. Training at 0 ms leads to limited generalization to post or prior contextual movement timing relationships.

The temporal generalization of the learned motor memory was again examined at the end of the second day of the experiment, using probe trials at different contextual timing relations ranging between -1.8 and +1.8 s. The temporal generalization of the motor memory after training at 0 ms had a very strong decline in compensation when the onset of the contextual movement was either advanced or delayed (Fig. 3G). Post-hoc comparisons supported the significant decreased away from the trained condition. The simultaneous timing was significantly different than all post (all p<0.001) and all prior (all p<0.001) timings, but there were no significant differences between any of the post and prior timings (all p=1.0). This generalization pattern was consistently expressed by all participants (Fig. 3F). Overall, we find that while strong learning occurs in the 0 ms condition (50%), there is limited transfer to post contextual movements (<10%), and no transfer to prior contextual movements (0%).

### Experiment 3: Prior Adaptation Context

In the third experiment (prior context), right-hand contextual movements were conducted prior to left-hand adaptation movements (-750 ms). During the pre-exposure phase, participants made straight reaching movements (Fig. 4A), which were strongly disturbed in the initial exposure of the force field (Fig. 4B). However, by the end of the exposure phase the participants made fairly straight movements to the targets (Fig. 4C) indicating that some compensation to the applied dynamics was achieved. Again, the kinematic error (MPE) increased immediately upon introduction of the force field (Fig. 4D), but reduced over the course of force field exposure. A repeated measures ANOVA found significant differences across the phases of the experiments (F_2,14_=228.999; p<0.001; ω^2^=0.944), with initial exposure larger than pre-exposure (p<0.001) and final exposure (p<0.001), although final exposure was still higher than pre-exposure (p<0.001). Force compensation increased throughout the exposure phase, reaching approximately 40% by the end of the second day (Fig. 4E). A repeated measures ANOVA found a significant increase in force compensation between final pre-exposure and final exposure levels (F_1,7_=67.810; p<0.001; ω^2^=0.818).

**Figure 4:**
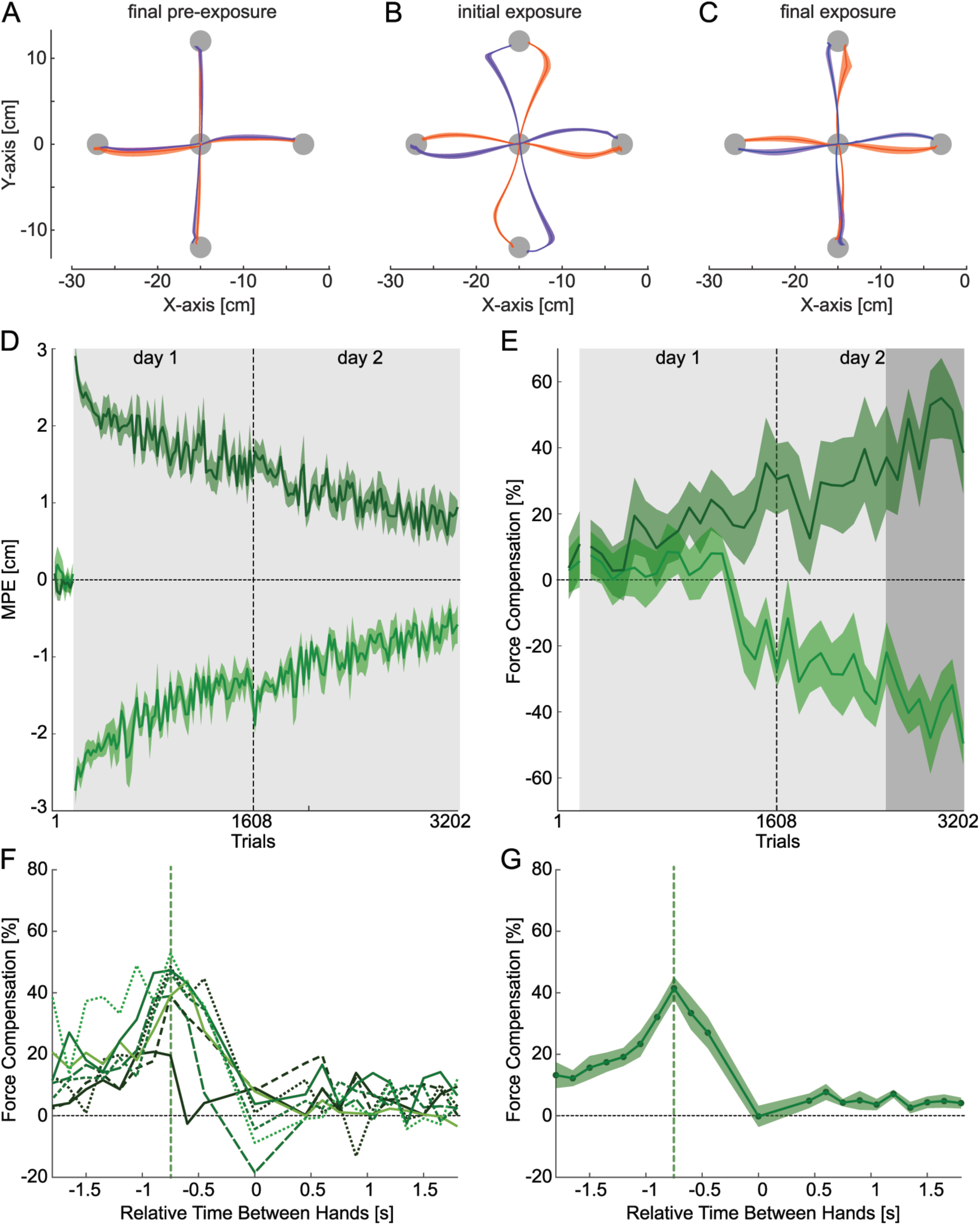
Results for Experiment 3 where the left-hand adaptation movement followed the right-hand contextual movement by 750 ms (-750 ms condition). **A.** Hand paths in the last block of pre-exposure in the null field condition. **B.** The hand paths on initial exposure to the curl field. **C.** The hand paths in the final block of curl field exposure. **D.** The kinematic error (MPE) for the left-hand for each curl force field direction separately. **E.** Force compensation for each curl force field direction separately, expressed as a percentage of complete adaptation to the force field. **F.** The temporal generalization of the learned motor memory to all timing relations between the two hands shown for individual participants. Each line denotes a different participant and the green dashed vertical line shows the trained relation. **G.** The mean and standard error of the temporal generalization of the learned motor memory across participants. Training at -750ms leads to a swift decline in generalization to other prior adaptation contextual movement temporal relationships, with no generalization to post contextual movements.

The temporal generalization of the learned motor memory was once again examined at the end of the second day of the experiment, using probe trials at different contextual timing relations ranging between -1.8 and +1.8 s. The temporal generalization of the motor memory after training at -750 ms shows a marked decrease in compensation when the onset of the contextual movement deviated from the trained timing relation (Fig. 4G). The trained timing (-0.75 s) force compensation generalizes well to nearby timings (-0.45 s, -0.6 s, -0.9 s), with no significant differences between any of the three timings (all p>0.685). However, the force decreases away strongly from the three times with the strongest responses (-0.6 s, -0.75 s, and -0.9 s) to the 0 ms timing (all p<0.001), all post timings (all p<0.001), and all prior timings less than or equal to -1.6 s (all p<0.01). The decay to longer prior adaptation contextual movement times already decreased from the peak compensation by the 1.05 s timing (p=0.022). However, no significant differences were found between any of the post, simultaneous, or prior timings less than -1.35 s (all p>0.491). This generalization pattern was consistent across all participants (Fig. 4F). Overall, we observed a significant decrease in compensation when the onset of the contextual movement was either advanced or delayed relative to the training time, even for other prior adaptation contextual timing relations. There was no transfer to the synchronous timing relation or to prior movements.

### Comparison Across Experiments

The three experiments followed identical designs, except for the timing relation between the two arms in the training condition, allowing for comparison across experiments. At the end of the force field exposure (final block of day 2), the simultaneous 0ms condition (Experiment 2, blue) yielded the straightest left-hand trajectories (Fig. 3B), whereas both the post (Experiment 1, red) and prior (Experiment 3, green) contexts remained slightly more curved (Figs. 2B, 4B). These trends align with the final levels of MPE values observed at the end of the experiments (Fig. 5A, B). The simultaneous context (blue) exhibited the most rapid decrease in MPE, whereas the prior adaptation context (green) and post contexts (red) had slower reductions in MPE. However, all three experiments showed clear reductions in MPE over the course of the experiment. An ANOVA comparing the final levels of MPE in the last block of the experiment found significant differences between the experiments (F_2,21_=7.503; p=0.003; ω^2^=0.028), with the simultaneous context showing a lower final MPE than either the post (p=0.004) or prior (p=0.023) contexts, but no significant differences between the post and prior contexts (p=0.341).

**Figure 5.**
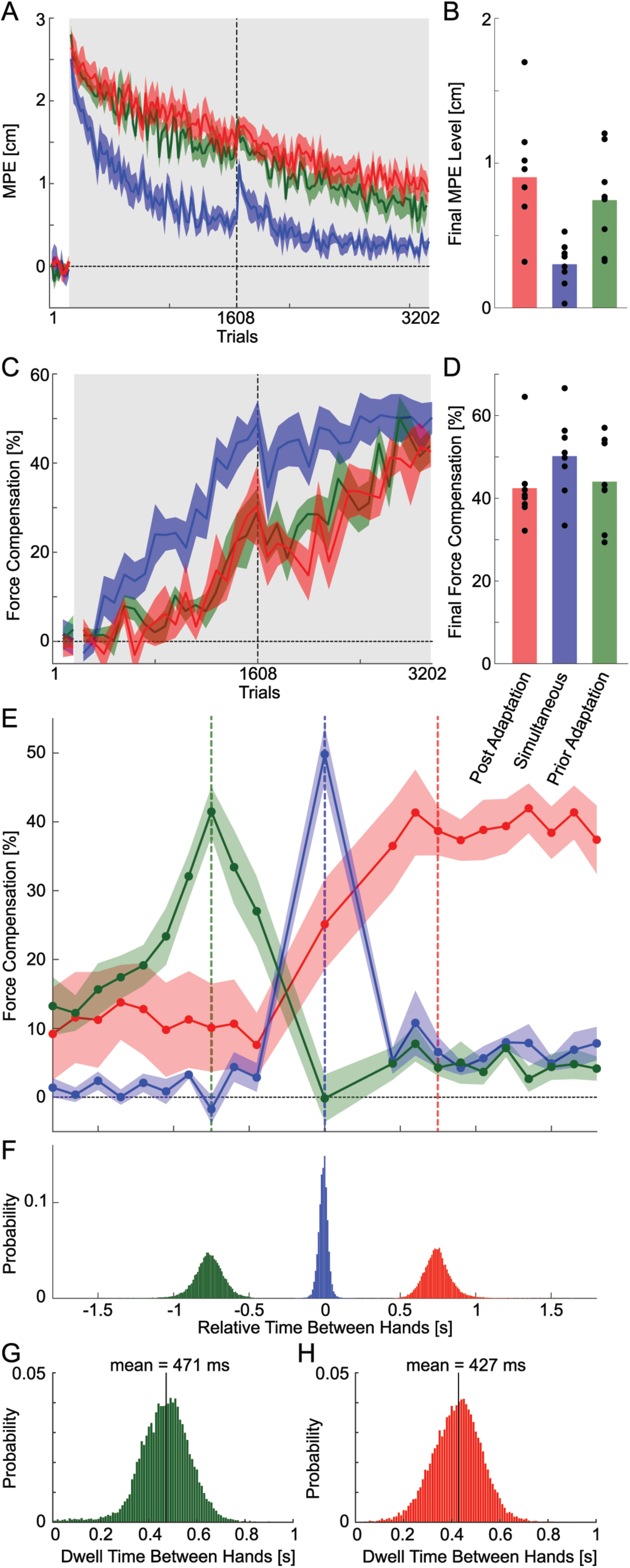
Comparison across the three experiments where participants were trained with either post (+750 ms; red), synchronous (0 ms; blue) or prior (-750 ms; green) timing relation. **A.** Comparison of kinematic error (MPE) changes across the experiment. **B.** Final levels of MPE at the end of the experiment. Colored bars indicate the mean MPE across participants in the final exposure block, while black circles represent individual participant values. **C.** Force compensation evolution across the three experiments. **D.** Final force compensation values across the three experiments (mean and individual participants) are shown for the final exposure block. **E.** Comparison of temporal generalization across bimanual timing relations after training at one of the three timing relations. Colored vertical lines illustrate the trained relation for each experiment. Solid lines depict the mean, and shared regions the standard error of the mean, across participants. Dot placements on the plot indicate the central location and the estimated mean compensation value for each bin. **F.** Histograms present relative movement initiation offsets between the two arms for all null practice and curl field training trials across the three experiments. Results for the +750 ms post movement context experiment are in red, the synchronous 0 ms movement context experiment in blue, and the -750 ms prior movement context experiment in green. **G.** Histograms of the dwell time in the prior adaptation movement context. **H.** Histograms of the dwell time in the post adaptation movement context.

Force compensation exhibited similar patterns across all three experiments (Fig. 5C). The simultaneous context (blue) showed the fastest increase in adaptation, with little differences in the development of the force compensation between the post (red) and prior (green) contexts. However, all three experiments achieved comparable final compensation levels at the end of the experiment (Fig. 5D), with no significant differences across the three conditions (F_2,21_=1.350; p=0.281; ω^2^=0.351). Overall, the results indicate that the timing relationships in all three contextual timing relationships act as effective cues for learning and adapting to opposing dynamics in the contralateral arm, facilitating the adaptation to opposing force fields.

A comparison of the temporal generalization across the three experiments reveals the distinct effects resulting from training at the three distinct timing relations (Fig. 5E). The synchronous context (blue) has the highest peak of adaptation at the training condition with negligible generalization to either post or prior timing relations. In contrast, both the post (red) and prior (green) timing training contexts demonstrate generalization across various timing relations. However, the prior context (green) shows selective generalization within different prior timings (decreasing away from the training condition), and no transfer to the synchronous or post timings. Conversely, the post context (red) exhibits strong generalization across all post timing relations, partial transfer to the synchronous condition, and limited transfer (10%) to prior conditions. Post-hoc comparisons showed significant differences between the three experimental conditions, with the post context (red) exhibiting significant differences (all p<0.001) to both the simultaneous and prior contexts at every post timing (+1.8 s to +0.45 s). All three contexts were significantly different from one another at the simultaneous 0.0 s timing (all p<0.03). Similarly, the prior context was different from both other contexts at the trained 0.7 s timing (all p<0.001). The prior context (green) was also significantly different from the simultaneous context (blue) over all tested timings between 0 s and -0.9 s (all p<0.043). However, there were no significant differences between any of the three contexts in the timings from -1.06 to -1.8 s (all p>0.123). Overall, we find distinct temporal generalization across the three experiments.

To perform a similar comparison but without binning the data, we additionally used generalized Additive Mixed Models (GAMM). The GAMM analyses demonstrated that the effect of time on the force compensation significantly varies across conditions, as indicated by a likelihood ratio test comparing the full model (which allowed trends to differ by condition) to the reduced model (which assumed a common trend) (χ^2^(4)=1009.33, p<0.0001). Subsequent pairwise comparisons confirmed that the temporal force trends differ significantly between each pair of conditions (all p<0.0001), highlighting the distinct temporal dynamics associated with each condition.

We propose that the variations in the temporal generalization patterns across the three experiments are indicative of different tuning functions within the sensorimotor control system for contextual movements. These variations may play a key role in the learning of coordinated actions across limbs. However, an alternative explanation could be different levels of variability in the training times, which might directly contribute to these patterns. To explore this possibility, we quantified the variability in training timing for each experiment (Fig. 5F). Here each histogram represents the distribution of the relative start times between the arms during the training movements for each experiment. The synchronous context (blue) had the narrowest spread of timings in the training condition, with times very close to the requested 0 ms separation. In contrast, the training conditions for the post (red) and prior movement adaptation (green) contexts exhibited slightly greater variability in onset times. However, this variability was significantly smaller than the width of the generalization tuning functions observed, particularly in the case of the post and prior contexts. This implies that the observed generalization is more likely to reflect underlying neural circuitry rather than being merely a result of variations in the training conditions.

To further quantify movement timing relationships, for Experiments 1 and 3 we computed time differences histograms between the timing between the start of the contextual movements after the end of the adaptation movements, and between the end of the contextual movements and start of the adaptation movements respectively (Figs. 5G, H). These values reflect the dwell time between the subsequent movements, with mean values of these distributions of 427 ms and 471 ms respectively.

## Discussion

We examined the relationship between right arm contextual movements and motor learning and recall in the left arm using a force field interference task. In three experiments, participants simultaneously adapted to two opposing force fields, where the force field directions were uniquely associated with the movement angle between arms. Each experiment trained participants at a distinct timing relation between the two arm movements (prior, simultaneous, or post-context) and assessed temporal generalization. Motor memory generalization across the experiments yielded three key findings. First, training in the simultaneous context led to recall primarily only at the learned timing, with limited transfer. Second, post-adaptation movement training resulted in broad generalization to other post timings and to a lesser extent the simultaneous timing. Lastly, prior-adaptation context training led to generalization to similar prior timings, and no significant transfer to simultaneous or post contexts. Overall, we find distinct patterns of temporal generalization based on the training, emphasizing the unique characteristics and implications of prior, simultaneous, and post-adaptation contexts in bimanual dual adaptation.

The results confirm that bimanual motion is a strong contextual cue for learning opposing dynamics. The relative motion of one limb predictively influences forces in the other limb, regardless of timing. This aligns with research showing that simultaneous movements reduce interference during dual adaptation (Nozaki et al., 2006; Howard et al., 2010; Yokoi et al., 2011). For example, switching between motor memories is facilitated when one hand is moving or stationary (Nozaki et al., 2006), when the limbs move in the same or opposite directions (Howard et al., 2010), or when the relative angle between hands varies (Yokoi et al., 2011).

In unimanual movements, prior (Howard et al., 2012, 2020; Howard and Franklin, 2015) and post adaptation movements (Howard et al., 2015; Sheahan et al., 2016), facilitate learning separate motor memories. Prior motion also influences learning in bimanual movements (Gippert et al., 2023). Our study demonstrates that post adaptation contextual movements also drive learning of independent motor memories in the contralateral limb, similar to unimanual studies (Howard et al., 2012, 2015; Sheahan et al., 2016). While we found comparable adaptation levels in both prior and post adaptation contexts, unimanual studies showed almost double compensation for prior (lead-in) compared to post-adaptation (follow-through) movements.

In unimanual dual adaptation, prior contextual movements are most effective when they directly precede adaptation movements, with the contextual effect diminishing as the dwell time (the time between the end of the contextual lead-in movement and the start of the adaptation movement) increases (Howard et al., 2012). In Experiment 3 the dwell time of 471 ms produced about 40% force compensation. Although this dwell time produced similar levels of adaptation in lead-in unimanual movements (Howard et al., 2012), higher adaptation occurred for shorter dwell times. In Experiment 1 the dwell time of 421 ms produced 40% adaptation, similar to that for follow-through unimanual movements (40%) with dwell times close to zero (Howard et al., 2015).

One key difference in our experimental design to previous bimanual studies (Nozaki et al., 2006; Howard et al., 2010; Yokoi et al., 2011; Gippert et al., 2023), is that our participants never received online visual feedback about the movement of their contextual (right) arm, whereas such feedback was provided in the previous research. This control is crucial because visual cursor motion alone is a strong contextual cue (Howard et al., 2012, 2020; Howard and Franklin, 2016; Franklin et al., 2023). Therefore, the inclusion of visual motion in previous bimanual studies might reflect the impact of this visual element rather than the physical motion of the contralateral limb. Although Gippert and colleagues (Gippert et al., 2023) included a visual ‘bimanual’ condition showing little adaptation, the visual cursor was offset by 10 cm from the adaptation movement. This offset might have weakened the contextual strength or acted as a weaker peripheral visual cue (Howard et al., 2013). Our study demonstrates that prior, simultaneous, and post adaptation movements of the contralateral limb can facilitate the separate learning of motor memories, even without visual information. This finding supports the interpretations of previous studies (Nozaki et al., 2006; Howard et al., 2010; Yokoi et al., 2011), and highlights the robustness of limb movement as a contextual cue in motor learning.

Numerous studies have demonstrated generalization of a motor memory across movement directions (Thoroughman and Shadmehr, 2000; Donchin et al., 2003; Gonzalez Castro et al., 2011), kinematics (Joiner et al., 2011; Howard et al., 2020; Orschiedt and Franklin, 2023) and postures (Shadmehr and Mussa-Ivaldi, 1994; Berniker et al., 2014; Leib and Franklin, 2021). Our study is the first to show temporal generalization. Just as generalization across directions has been used to infer the neural basis functions underlying motor memories (Poggio and Bizzi, 2004; Shadmehr, 2004; Joiner et al., 2011; Kadiallah et al., 2012), we propose that the pattern of the temporal generalization reflects the temporal tuning of these neural basis functions.

The three experiments exhibit different levels of variability in the timing between limbs during training (Fig. 5F). While the prior and post adaptation movement contexts show similar timing variability, the simultaneous context has a narrower range, indicating greater consistency. This is unsurprising as simultaneous bimanual movements are generally easier to perform (Kelso, 1995; Howard et al., 2009a; Kobayashi and Nozaki, 2023). Kinematic variability of contextual movements during adaptation reduces the learning rate (Howard et al., 2015, 2017). Therefore, temporal variability of contextual movements likely had a similar effect, as we observed slower final adaptation levels for the prior and post contexts. Our study is unable to separate the effect of temporal variability from the training condition. The prior and post conditions exhibit broader temporal generalization than the simultaneous condition, which shows almost no generalization to other timings. Despite similar training time variability, strong differences in generalization between prior and post contexts exist. The post adaptation context generalizes to simultaneous and prior adaptation times, suggesting generalization is not confined to training variability. Both prior and post contexts show generalization beyond training variability. We propose that the observed differences stem from variations in the inherent use of contextual information for generating natural behavior, and therefore in the underlying neural circuitry involved in learning and executing coordinated bimanual movements.

Neural activity associated with generating and controlling bimanual movement (Swinnen, 2002) involves interactions between both cerebral hemispheres (Gerloff and Andres, 2002). This includes both contralateral and ipsilateral primary motor cortex, where bimanual and unimanual movements exhibit different neural activity patterns (Donchin et al., 1998; Steinberg et al., 2002; Cross et al., 2020). The cerebellum (Wolpert et al., 1998; Imamizu et al., 2000; Ito, 2005), parietal, and frontal areas also play significant roles (Scott, 2012). While we cannot elaborate on the neural regions responsible, our work examines the temporal tuning of the neural basis functions that lead to motor memories through generalization. Local neural basis functions are suggested to underlie motor memory formation (Poggio and Bizzi, 2004; Shadmehr, 2004; Joiner et al., 2011; Kadiallah et al., 2012), allowing generalization to similar states. These basis functions often exhibit Gaussian-like tuning. However, our findings reveal that temporal generalization can exhibit different forms depending on the training conditions, ranging from broad temporal generalization in the prior context to narrow tuning in the simultaneous context.

A normative perspective can provide insights into the human nervous system’s operation (Franklin et al., 2016; Makino et al., 2016; Česonis and Franklin, 2022; Leib et al., 2024). Prior movements are crucial for anticipatory muscle activation patterns in real-world tasks (Klapp, 1979), especially in bimanual tasks, where coordination and anticipatory adjustments between limbs ensure synchronized actions (Swinnen and Wenderoth, 2004). As actions at one time can require compensatory actions at different times, this might explain why both prior and past movements show strong temporal generalization. Adjustments to forces produced by the other limb are relevant in simultaneous movements, such as opening a jar or pouring a glass of water. However, these interactions are so closely linked in time that are often taken as evidence for predictive models (Desmurget and Grafton, 2000; Wolpert and Flanagan, 2001; Flanagan et al., 2003; Franklin and Wolpert, 2011). This might explain why trained simultaneous interactions show limited generalization to prior or post-adaptation contexts.

Considering an evolutionary perspective is useful (Cisek, 2022; Cisek and Hayden, 2022). In locomotion on variable ground, movement of one limb requires compensatory motions in the other limb, both for prior movements and in preparation for future steps. The temporal interval between steps can vary widely, which might explain the plastic adaptation and generalization to prior and post-adaptation movements, while narrow generalization of simultaneous movements reflects tight coupling in object manipulation.

This work highlights the sensorimotor control system’s plasticity, showing different training conditions produce widely different temporal generalization. These differences might be explained by considering the influence of different tasks on the neural circuitry over evolutionary timescales. While state-space (Lee and Schweighofer, 2009; Forano and Franklin, 2020), and contextual inference models (Heald et al., 2021, 2023) can explain dual-adaptation, these models cannot yet predict different patterns of temporal generalization.

## Acknowledgements

Financial support was provided by the Technical University of Munich and the SECAM at the University of Plymouth. During the preparation of this work, ISH utilized ChatGPT-4o for proofreading. We thank Yiming Liu for assistance with the Generalized Additive Mixed Models.

## Author Contributions

The study was conceived and designed by ISH and DWF. The study was implemented by ISH. The experiments were conducted by SF. Data analysis was carried out by ISH. Figures were prepared by ISH and DWF. Statistics were performed by DWF. The manuscript was written by ISH, SF and DWF. The final manuscript was reviewed and edited by ISH, SF and DWF.

## Notes

Statement Financial interests or conflicts of interest: The authors declare that they have no financial, personal, or professional interests that could be construed to have influenced the paper.

### Competing Interest Statement

The authors have declared no competing interest.

### Summary of Updates

We have made a few minor changes to the manuscript

